# Structured Inhibitory Activity Dynamics During Learning

**DOI:** 10.1101/566257

**Authors:** Moises Arriaga, Edward B. Han

**Affiliations:** Department of Neuroscience, Washington University School of Medicine, St. Louis, MO, USA

## Abstract

Inhibition plays a powerful role in regulating network excitation and plasticity; however, the relationship between learning and inhibitory activity dynamics remains unclear. Furthermore, do individual interneurons play consistent functional roles during learning or is their activity driven stochastically? Using two-photon calcium imaging, we recorded hippocampal CA1 somatostatin-expressing interneurons as mice learned a virtual reality task in new visual contexts. We found strong initial suppression of inhibitory activity that gradually diminished as animals learned, while blocking learning using a “no task” environment prevented the recovery of inhibitory activity. Although the magnitude of suppression differed across the population, each interneuron showed consistent levels of activity modulation during learning across multiple novel environments. This work reveals learning-related dynamic inhibition suppression, during which inhibitory networks display stable and consistent activity structure. This functional inhibitory circuit architecture suggests that individual interneurons play specialized and stereotyped roles during learning, perhaps by differentially regulating excitatory neuron ensembles.

## Introduction

Excitation is balanced by inhibition in neuronal networks (Andersen et al., 1963). In cortical circuits, feedforward inhibition is rapidly and robustly recruited by excitatory inputs, while pyramidal neuron firing elicits feedback inhibition to further dampen excitability (Alle et al., 2001; Lamsa et al., 2005; Pouille and Scanziani, 2001, 2004). Furthermore, inhibition strongly controls synaptic plasticity, a putative cellular mechanism of learning (Bliss and Lømo, 1973; Whitlock et al., 2006). Intact inhibition limits potentiation to relatively low levels while pharmacologically blocking inhibition facilitates both the induction and magnitude of potentiation (Artola and Singer, 1987; Bear et al., 1992; Steward et al., 1990; Wigström and Gustafsson, 1983). Thus, inhibition suppression is a potential mechanism for enhancing learning by favoring synaptic plasticity in excitatory neurons.

Notably, inhibition can be strongly modulated *in vivo* in freely moving rodents. CA1 fast-spiking putative interneurons are suppressed when rats explore a novel spatial environment, although these studies did not explicitly address task learning, as animals were foraging for random rewards (Frank et al., 2004; Nitz and McNaughton, 2004; Wilson and McNaughton, 1993). Learning new food locations in a familiar environment dynamically modulated fast-spiking interneuron activity and altered their associations with pyramidal cell ensembles (Dupret et al., 2013). Hippocampal CA1 parvalbumin-expressing interneurons (PV-ints) have decreased network connectivity during initial learning of the Morris Water Maze that then increases with task performance and this modulation is necessary for learning (Donato et al., 2013; Ruediger et al., 2011). Furthermore numerous studies have shown suppression of SOM-and/or PV-ints is necessary for certain types of cortical and amygdalar learning, often triggered by disinhibitory inputs from interneurons that preferentially target other interneurons for inhibition, including vasoactive intestinal peptide-expressing interneurons (VIP-ints) (Gentet et al., 2012; Karnani et al., 2016; Lee et al., 2013; Letzkus et al., 2011; Mardinly et al., 2016; Pi et al., 2013; Turi et al., 2019; Wolff et al., 2014). Finally the activity or plasticity of numerous other interneuron cell-types have been implicated controlling animal behaviors (Basu et al., 2016; Hartzell et al., 2018). Together, this work demonstrates the dynamic nature of inhibition during learning and identifies the importance of inhibition suppression in certain types of learning.

Our understanding of *in vivo* inhibitory activity in the hippocampus is primarily driven by recordings of soma-targeting fast-spiking interneurons (likely PV-ints) since their distinctive firing characteristics make them identifiable in extracellular electrophysiology recordings. SOM-ints, while far less understood, are particularly interesting because they selectively innervate the dendrites of pyramidal neurons and can directly control dendritic excitability. Dendritic spikes, typically characterized by calcium entry through Ca^2+^ channels or NMDA receptors, generate burst firing of neurons and can mediate long-term plasticity, place field formation, and learning (Bittner et al., 2015, 2017; Cichon and Gan, 2015; Larkum et al., 1999). Formation of place fields is associated with dendritic spikes that occur during transient periods of SOM-int activity suppression in novel environments (Sheffield et al., 2017). However, there is less evidence for SOM-int suppression during learning in the hippocampal formation. SOM-int activation rather than suppression is required for fear learning, both in CA1 or the dentate gyrus (Lovett-Barron et al., 2014; Stefanelli et al., 2016). The activity dynamics of SOM-ints during more complex learning paradigms remains unknown.

While interneuron activity can be dynamically modulated during learning, the stability of these activity dynamics across learning episodes has not been studied. This is partially a technical issue since extracellular electrode recordings are typically stable for a few hours and have limited ability to identify interneurons, making longitudinal recording of single interneuron activity difficult. Conceptually, interneuronal activity is thought to be governed by pyramidal neurons, driven in a feedforward and/or feedback manner. Thus, if activated pyramidal neuron ensembles are stochastic, stochastic ensembles of interneurons should result. An alternate possibility is that individual interneurons play consistent and reproducible roles during learning, reflecting an underlying structure in how inhibition regulates the network. While inhibitory neurons are composed of multiple cell-types playing distinct network roles (Klausberger and Somogyi, 2008; Pelkey et al., 2017; Wamsley and Fishell, 2017), little is understood about the functional specialization of interneurons within a defined cell-type (Arriaga and Han, 2017).

Here we examined the structure of SOM-int inhibition by calcium imaging over multiple days and learning episodes to determine both the learning-related changes and the consistency of individual interneurons. We found SOM-int activity was strongly suppressed in novel environments, with activity gradually returning as animals re-learned goal locations. In contrast, for animals in an environment with no relevant learning (static visual scene with no task), inhibition remained persistently suppressed over days, suggesting that recovery of inhibition is tied to learning rather than habituation. Surprisingly, suppressed inhibition triggered by context changes showed a defined population structure during both learning in the novel environment and the “no task” condition. Each interneuron exhibited consistent activity suppression, with high correlation of suppression across multiple novel contexts as well as the “no task” condition. These data reveal interneuron activity suppression during remapping and learning in the hippocampus and reveal functional inhibitory structure that may route the encoding of information within the pyramidal network.

## Results

### Virtual reality (VR) behavior and learning

To study the activity dynamics of SOM-ints during learning, we used two-photon calcium imaging to stably record from neurons over weeks. Our goal was to study the activity dynamics of the same cells in a goal-directed, spatial learning task. We initially trained water scheduled mice to run to alternating ends of a virtual visual track to receive water rewards, using their movement on a floating spherical treadmill (styrofoam ball) to control their movement in VR (Arriaga and Han, 2017; Dombeck et al., 2010; Harvey et al., 2009). Mice need to physically rotate the ball at the ends of the track to turn around in VR and run forward on the ball to reach the opposite end.

We modified this task to study learning by exposing animals to a novel VR environment where mice need to re-learn the task with new visual cues. We ran animals for 7 min in the well-trained familiar environment (Fam), immediately switched mice to a new VR environment for 14 min (New), and then back for another 7 min session in the original familiar environment (Fam’) (schematic, Figure 1A). The task was identical in familiar and New epochs but the visual textures of the walls and tracks change, as well as positions and textures of distal landmarks surrounding the track. We repeated this protocol over five days, with the same New environment each time. This paradigm has numerous advantages for studying learning. First animals are already well-trained in the task in the familiar environment, which allows mice to learn at a much faster rate in a novel context (2–5 days to achieve high behavioral performance) than the initial learning of the task (2 – 5 weeks). Second, the immediate switch from a familiar to a novel context marks a highly salient and defined start to the learning period. Finally, a return to the familiar environment (Fam’) allows us to verify that impaired behavioral performance in New is not due to time-dependent effects such as satiation or fatigue.

**Figure 1:**
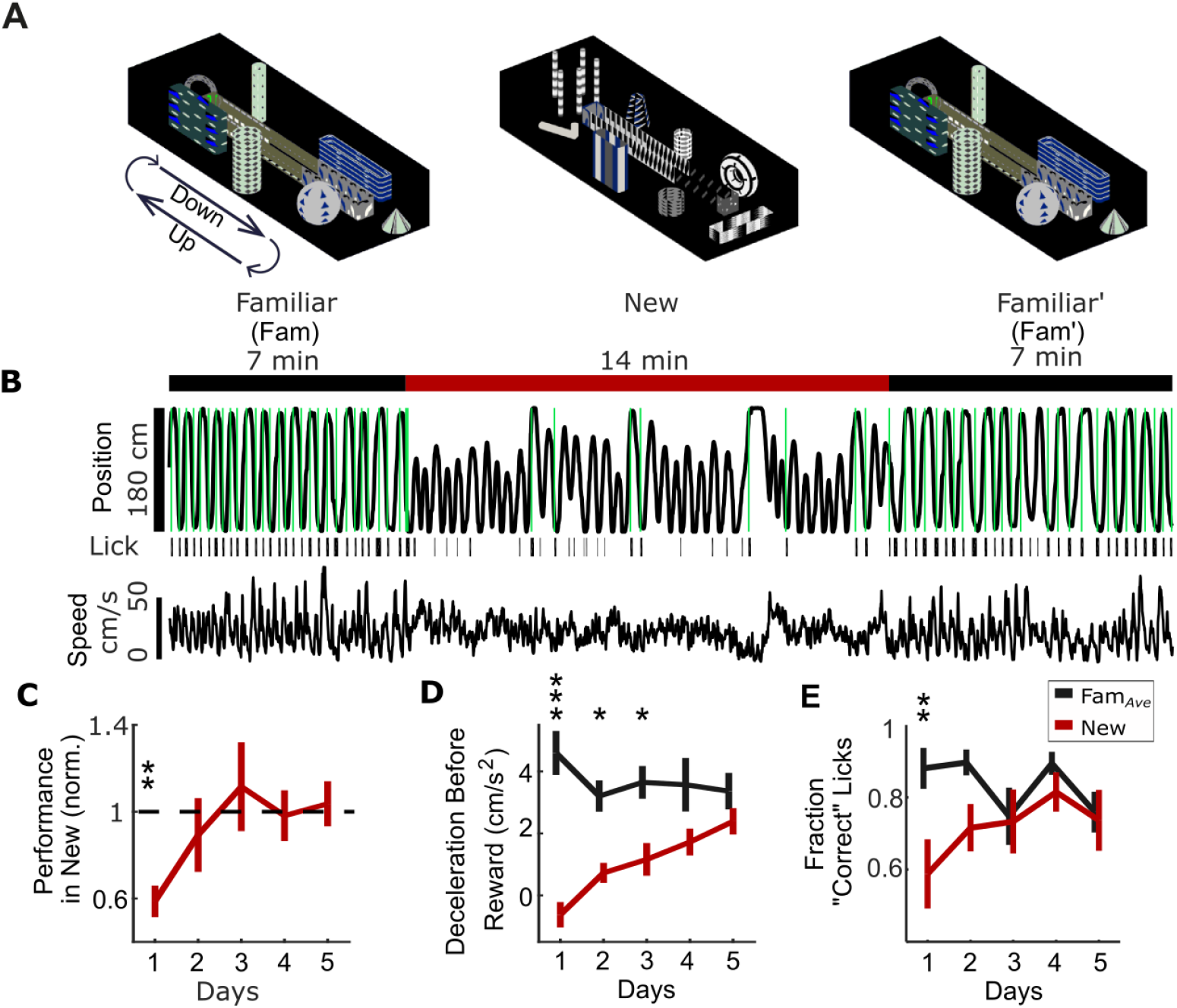
Learning in new visual virtual reality (VR) environments. **A**, Head-fixed mice run to alternating ends of the VR track by controlling movement of a floating spherical treadmill (Styrofoam ball). Mice run forward on the ball to traverse the track and rotate the ball to turn around in VR. Animals spend 7 min in a familiar environment (Fam), which is instantaneously replaced with a new environment (New) for 14 min, before returning to the same familiar environment (Fam’). The task is the same but the visual scene differs across environments. **B**, Example mouse position in VR shows running to alternating ends of track with water rewards (green) in Fam., with worse performance in New. Lick bouts (black bars) are tracked with an electronic sensor on the lick tube. Ball speed shows similar magnitude in New and Fam. environments. **C**, Behavioral performance is initially impaired in New (rewards/min in New normalized to Fam_*Ave*_, the average performance in flanking Fam and Fam’ epochs) but improves over time. **D**, Mice slow down prior to reward in the familiar environments, measured as deceleration in the 3s window before reward. Deceleration before reward is initially lower in New but increases over days, suggesting learning of reward sites. **E**, Most licks in Fam_*Ave*_ are within a 1s window centered on reward delivery (defined as “correct” Licks). Fraction of “correct” licks is initially lower in New relative to Fam. but increases over days, again suggesting learning of reward sites in New (*p<0.05, **p<.01, ***p<.001 by paired sample t-test or 1-sample t-test with Bonferroni-Holm Correction, N=9).

To better understand the behavioral effects of the familiar to novel environment switch and investigate links to learning in New, we characterized behavioral performance using a large cohort of mice, a subset of which were used for SOM-int imaging (the remainder were used for other experiments and not discussed here). We quantified task performance as number of rewards per minute (rew/min) in Fam, New, and Fam’ epochs. Upon initial exposure to the New virtual world, animal behavior was dramatically altered. Performance in New was significantly worse on day 1, compared to Fam_*Ave*_ (the average performance in the flanking Fam and Fam’ epochs) (Figure 1C, performance in New normalized to Fam_*Ave*_ on day 1=0.59±0.07, p<0.001). This impairment was largest on day 1 and gradually decreased over the next four days of exposure to the same “new” world (Spearman ρ=0.41, p<0.001).

If these performance increases indicate learning in the New environment, then animals should also exhibit signs of increased familiarity with goal locations and expectations. In familiar environments, animals typically slow down before entering the reward zone (marked by a period of deceleration several second prior to the reward), in anticipation of receiving reward and consuming water (Gauthier and Tank, 2018). In new environments, animals initially did not decelerate as they approached the reward zone, but over 5 days of repeated exposure to New, they decelerated more and more, approaching Fam_*Ave*_ levels (Figure 1D, Day 1, p<0.01, Day 2 and 3, p<0.05). Similarly, animals in familiar environments generally begin licking within a 1s window centered on reward delivery (which we define as “correct” licks); in New, animals initially lick outside of this window (Figure 1E, Day 1, p<0.01), with their behavior improving over repeated training. These metrics support the interpretation that progressive improvements in rew/min in the New environment are due to animal learning. We characterized several other metrics of learning (Figure 1—figure supplement 1), supporting the notion that animals are engaged in day-over-day learning in the New environment.

### Characterization of neuronal calcium activity in novel virtual environments

To investigate *in vivo* interneuronal activity dynamics during exposure to novel environments, we used two-photon imaging of neuronal calcium activity during a spatial navigation task in visual virtual reality (VR) (Arriaga and Han, 2017). We used an electric tunable lens to image a 3-D volume of mouse hippocampal CA1 by capturing sequential imaging frames along the z-axis moving from ventral to dorsal. Images captured neuronal somata from *stratum pyramidale* and *oriens*, over four to six planes at a frame rate of 5.2 – 7.8 Hz per plane. Cre-dependent AAV1-Syn-Flex-GCaMP6f was injected into SOM-cre^+^ transgenic mice to drive a genetically encoded calcium sensor specifically in SOM^+^ hippocampal interneurons. Calcium activity can be taken as a proxy for neuronal activity as multiple studies using simultaneous *in vivo* imaging and cell-attached patch electrophysiology on the same neurons have found strong correlation between spiking and calcium signals (Chen et al., 2013; Dana et al., 2016). We, and others, have measured the activity dynamics and coding properties of hippocampal neurons in visual virtual environments and found significant similarity between VR and real world behavior in several aspects of function such as place coding, direction-specificity, place cell remapping in novel environments, and interneuron activity correlation and anti-correlation with locomotion (Arriaga and Han, 2017; Gauthier and Tank, 2018; Hainmueller and Bartos, 2018; Harvey et al., 2009; Sheffield et al., 2017).

### SOM-int activity suppression during learning in new environment

To investigate the functional activity dynamics of SOM-ints during learning, we recorded calcium activity from the same cells as animals performed the VR track running task in Fam, New, and Fam’ over five days. SOM-int neuronal activity, measured as ΔF/F, was strongly suppressed upon transition into New (ΔF/F of 6 sample cells from one imaging plane of one mouse, Figure 2A; Video 1 shows activity suppression in another set of SOM-ints). Individual neurons were differentially suppressed with some relatively unaffected. On returning to Familiar after New in Fam’, calcium activity rapidly recovered, as did behavioral performance. Similar results can be seen in all cells from this animal (Figure 2B). We quantified neuronal activity for all cells in this example animal as mean ΔF/F and compared across Fam, New, and Fam’. Activity suppression is calculated as the percent difference of mean ΔF/F for each cell between New and Fam_*Ave*_ using the formula 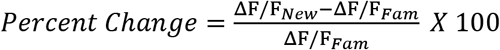. The histogram of percent differences for each cell in the sample mouse shows a distribution of cells that are suppressed in New (Figure 2C). The calcium activity from another sample mouse over five days exposure to New follows a similar pattern of activity dynamics (Figure 2—figure supplement 1). Across all animals, suppression histograms of SOM activity over the five day protocol show a large initial suppression in New that diminishes over days of exposure (Figure 2D).

**Figure 2:**
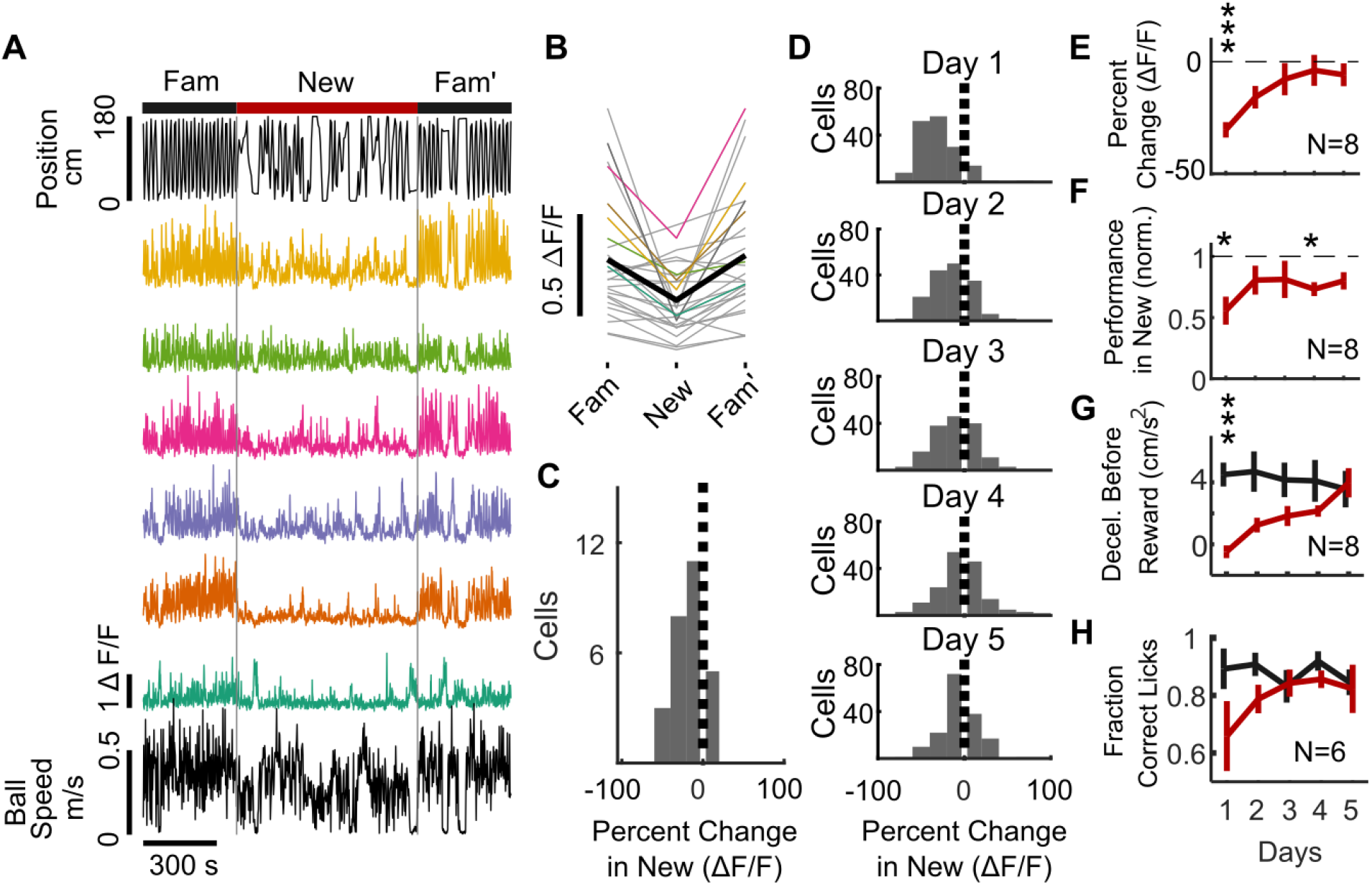
SOM^+^ interneuron (SOM-int) activity suppression in new environments. **A – C**, example data from individual mouse. **A**, Top, position in VR track of example mouse. Middle, ΔF/F of sample SOM-ints showing activity suppression in New. **B**, ΔF/F of all recorded cells (n=28 cells) from example mouse on day 1 of New exposure (colors) with mean of all cells (black). **C**, Histogram of percent change in ΔF/F of SOM-ints shown in (B), in New relative to Fam_*Ave*_ on day 1. **D**, Activity suppression in New decreases with exposure over days (N=8 mice, n=162 cells). **E**, SOM-int activity is initially suppressed but recovers over days of exposure to New (Average SOM-int activity per mouse). **F**, Performance in New world increases over days. **G**, Mice increasingly slow down prior to a reward suggesting re-learning of goal location in New. (N=8) **H**, No significant difference in “Correct” licks over days (note that N=6). (n.s. p>0.05, *p<0.05, **p<.01, ***p<.001 by paired sample t-test or 1-sample t-test with Bonferroni-Holm Corrections).

To quantify this suppression over time, we calculated a percent difference for each mouse by averaging all cells from each mouse and then calculated a grand mean for all mice on each day (Figure 2E). Indeed, SOM-int activity exhibited significant suppression in New that gradually decreased over days (mean suppression on Day 1=-30.41±3.46%, p<0.001). This decrease in suppression paralleled the increase in behavioral performance in New, as normalized to the average performance in Fam and Fam’ (Figure 2F, mean normalized performance on Day 1=0.56±0.11, p<0.05). (Note that behavioral metrics in Fig. 2 are from 8 SOM-cre imaged mice, a partially overlapping set of the larger behavioral cohort of 9 mice in Figure 1. Mice in Figure 1 all had lick sensor data, while not all mice in this group did). These data show, on average, strong suppression of SOM-int upon exposure to a New environment, with recovery of activity over repeated exposures (Spearman ρ=0.48, p<0.01). At the same time, behavioral performance was initially impaired and then increased over time (Figure 2G-H). These data are consistent with inhibition suppression acting as a permissive gate for learning; then, as performance improves, increased inhibition may serve to stabilize learning in the network.

A possible confound to this interpretation would be decreased locomotion in New. Many SOM-ints have activity that is positively correlated with locomotion, although there is a distinct population that is anti-correlated (Arriaga and Han, 2017). Thus, it is possible that decreased interneuron activity is due to decreased locomotion in New that gradually recovers over repeated exposure. While average locomotion was the same in New and Fam_*Ave*_ (Figure 1— figure supplement 1D), it remains possible that more nuanced changes in movement could result in decreased SOM-int activity.

To more thoroughly explore this possibility, we made a general linear model (GLM) to predict each cell’s fluorescence based on behavioral data. Models were trained using fluorescence data from Fam and fit using the forward and rotation components of ball speed, the timing of rewards, and position and speed in the VR environment. Modeled ΔF/F was very similar to actual ΔF/F in Fam, while in New, the fit of modeled ΔF/F was much worse (six sample cells, Figure 3A). To quantify this, we compared modeled ΔF/F to actual ΔF/F in two ways: root mean square (RMS) error, which decreases with better fit, and percent of variance explained (R^2^), which increases with better fit. Using both measures, we found model fit was significantly worse in New versus Fam_*Ave*_ for all five days suggesting that changes in behavioral variables, including locomotion, did not explain decreased ΔF/F in New (RMS error, Figure 3B; R^2^, Figure 3C). We also compared model fits in Fam, New, and Fam’ (rather than Fam_*Ave*_) and found that predicted cellular activity improved in Fam’ relative to New, although model fit in Fam’ was worse than in Fam (Figure 3—figure supplement 1). This could be due to residual activity suppression from exposure to the New world, or imaging-related photo-bleaching. Finally, to evaluate the contribution of individual behavioral variables to cellular activity, we quantified model fits based on single variables (Figure 3—figure supplement 2). Overall, models based on all variables had the best fits, while among the individual variables, those representing locomotion (forward and rotational ball speed, and VR speed) contributed the most to model fits of cellular fluorescence. These data show that the strong suppression of inhibitory activity in new environments is not simply explained by changes in mouse behavior but are likely to be triggered by contextual novelty itself.

**Figure 3:**
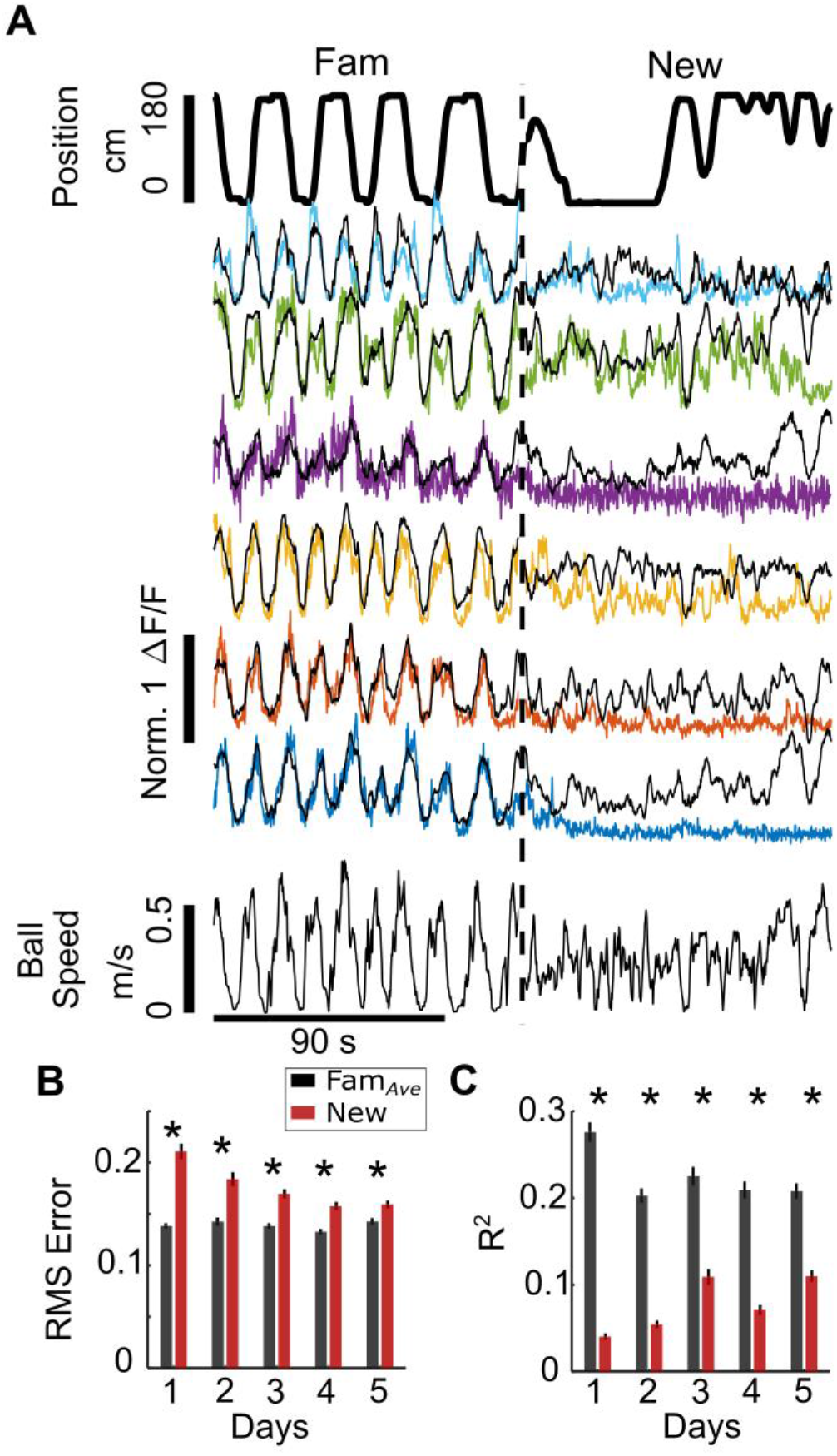
Decreased inhibition in New is not explained by altered behavior. **A**, Gaussian general linear models (GLMs) for individual SOM-ints were trained as a function of locomotion, VR movement, and rewards in Fam to predict calcium activity. In New, modeled ΔF/F (black) is larger than actual ΔF/F (colored traces), indicating that suppression of activity is greater than predicted from the model. **B**, Model fits are significantly worse in New versus Fam_*Ave*_ based on average Root Mean Square (RMS) error (lower errors mean better model fit). C, Average amount of variance (R^2^) predicted by model also shows worse model fit in New (greater R^2^ means better model fit) (*p<.001 by paired sample t-test Bonferroni-Holm Corrections, n=162).

Another potential interpretation of our data is that decreased SOM-int activity is driven by surprise at the context switch, with habituation to this surprise gradually restoring inhibitory activity, with no relationship between the return of inhibition and learning. To test this possibility, we dissociated surprise at the context switch from learning in the new environment by replacing the New environment with a no-task, no-reward epoch and a static visual scene (black screen). Under these conditions, if surprise drives inhibition suppression and activity recovery is due to habituation, we would see the same suppression and recovery over time as previously shown. On the other hand, if learning is necessary for the recovery of inhibitory activity and learning is prevented, we should see sustained inhibitory suppression.

Taking advantage of the long-term recording stability of two-photon calcium imaging, we recorded the same cells from a subset of the mice (N=4, n=69) used in the remapping experiment, allowing us to directly compare the kinetics of inhibition recovery when learning was present or absent. The “no task” environment evoked strong suppression of SOM-int activity, similar to the suppression seen when mice are switched into New (Figure 4 A-D, Mean percent change on Day 1=-48.10±4.18%, p<0.01). However, over five days of exposure to the same “no task” environment, inhibition remained strongly suppressed (Figure 4E, Figure 4—figure supplement 1, Spearman’s ρ=0.0188 p=0.939). In marked contrast, the same cells from the same mice showed strong recovery of inhibition over five days of exposure to New (Figure 4E, Spearman’s ρ=0.59, p<0.01). Thus, recovery of inhibition is unlikely to be the result of habituation to surprise and is more likely tied to learning in the new context.

**Figure 4:**
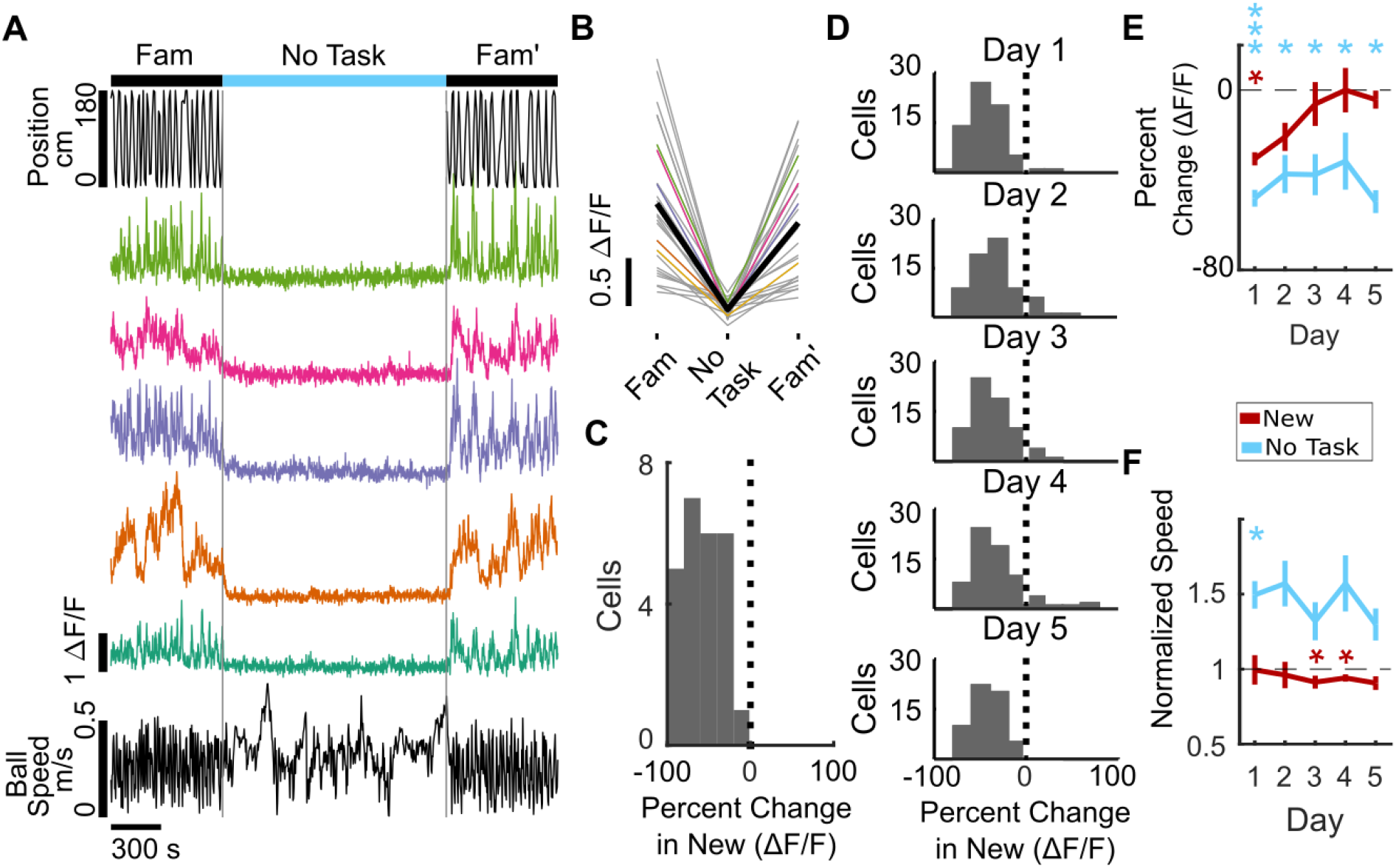
SOM-int activity suppression remains high when learning is blocked in “no task” environment. **A – C**, example data from individual mouse. **A**, Cells from sample mouse are strongly suppressed in “no task” epoch (static black screen, no rewards). Top, position in VR track, middle, ΔF/F of sample cells, bottom, ball speed. **B**, ΔF/F of all cells from example mouse on Day 1 of “no task” exposure showing activity suppression (mean of all cells in black). **C**, Histogram of percent change in ΔF/F of SOM-ints from example mouse on day 1 of “no task” showing strong suppression. (n=18). **D**, Interneurons remain suppressed over several days of “no task” exposure. Histogram of percent change of cells (N=4, n=69). **E**, In “no task” exposure, SOM-ints remain suppressed in contrast to recovery during exposure to New (from Figure 2E). **F**, Average speed in “no task” environment increases relative to Familiar, in contrast to same or decreased speed in New. (n.s. p>0.05, *p<0.05, **p<.01, ***p<.001 by paired sample t-test or 1-sample t-test with Bonferroni-Holm Corrections, N=6).

### Consistent inhibitory structure during learning

On average, SOM^+^ neurons are suppressed in a new environment (Fig. 2E), but the degree of activity suppression is heterogeneous across neurons, ranging from strong inhibition to moderate activation in individual cells (Figure 2D). This spectrum of interneuron activity modulation in novel environments could reflect changes in excitatory drive onto individual interneurons due to the stochastic activation of new place cell maps during global remapping in the hippocampus and grid cell realignment in the entorhinal cortex. Alternatively, it could indicate that different SOM-ints have distinct, but persistent, network roles. These two models have very different implications for interneuronal roles during learning. The first suggests a pooled inhibition model where each interneuron is “equi-potential” and has no particular specialized role in the network. The second signifies that network inhibition has a consistent structure, with the intriguing possibility that different interneurons play distinct functional roles.

We tested whether the structure of inhibition suppression was stochastic or consistent by putting a subset of the animals previously described through a second remapping protocol where they are exposed to another distinct and novel visual virtual environment, labeled “New 2,” with the original novel environment now labeled “New 1.” By recording the same cells across the two remapping protocols, we could correlate the magnitude of each cell’s activity suppression in New 1 vs. New 2. If SOM-ints are stochastically recruited by network activity, there should be no correlation in activity suppression across New 1 vs. New 2. However, we found strong correlation between activity suppression in New 1 vs. New 2 in individual SOM-ints. This correlation was strong on Day 1, and strikingly, this correlation was significant across all days of the remapping protocol (Figure 5A, Day 1, Pearson’s Correlation=0.70, p<0.001; Day 2, r=0.67, p<0.001; Day 3, r=0.76 p<0.001; Day 5, r=0.58 p<0.001). We, and others, have previously verified place cell global remapping across different virtual environments (Arriaga and Han, 2017; Gauthier and Tank, 2018; Hainmueller and Bartos, 2018; Sheffield et al., 2017), strongly suggesting that this consistent functional inhibitory network structure occurs despite differing ensembles of activated pyramidal neurons.

**Figure 5:**
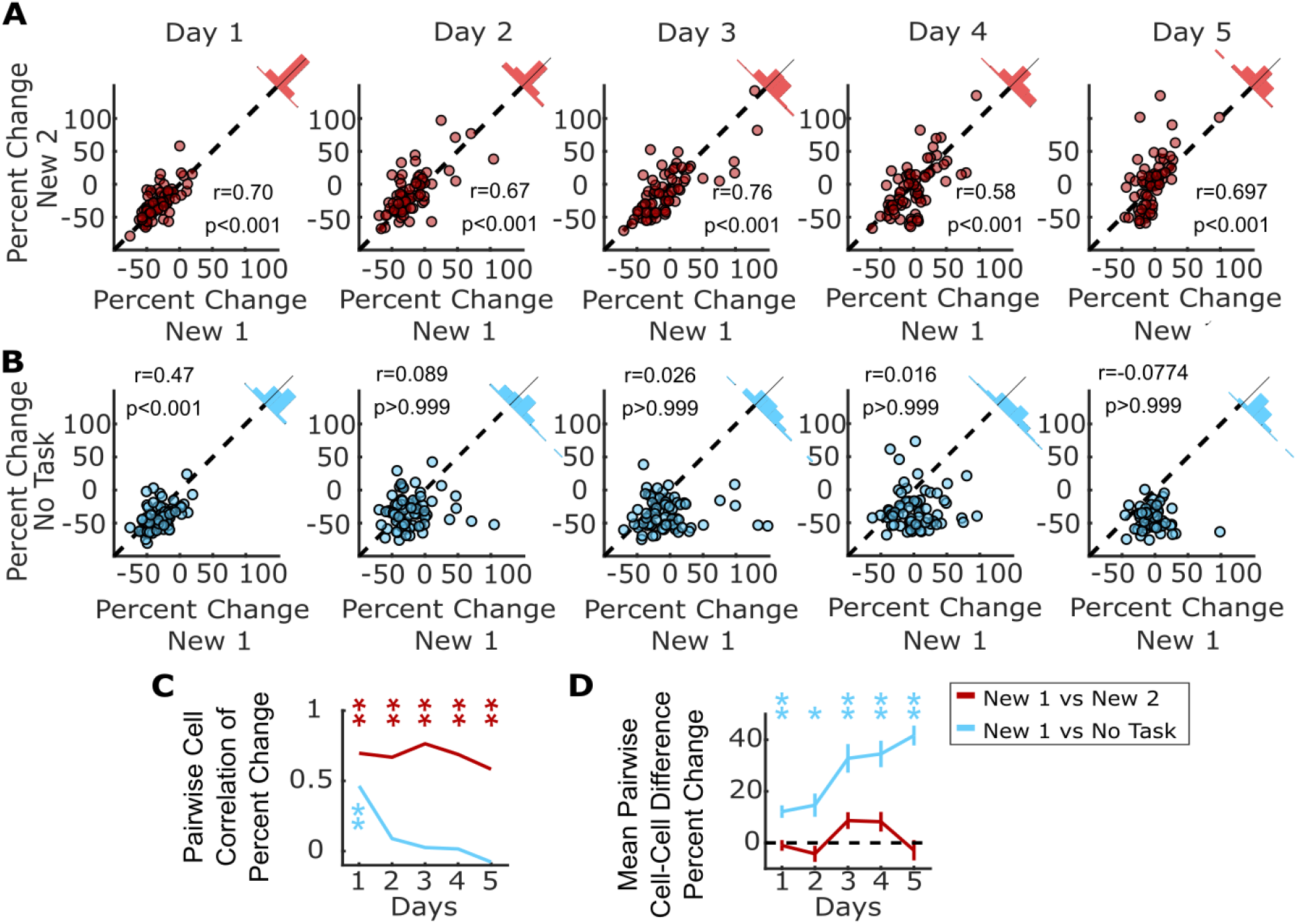
Consistent SOM-int activity responses across different new environments and in “no task” epoch. **A**, Individual SOM-ints show correlation in activity modulation in two distinct New environments. **B**, Similar correlation of activity modulation is seen between New 1 and “no task” exposures for Day 1. On subsequent days, correlation disappears as SOM-int activity begins to return in New 1 while remaining suppressed in “no task”. **C**, Summary of correlation data from (A) and (B). Correlation between percent change of cells between two remapping sessions or remapping session 1 and “no task” exposure session. **D**, Mean difference in percent change in activity in cells between remapping and “no task” exposure settings (*p<0.01, **p<.001 1-sample t-test with Bonferroni-Holm correction N=4, n=69).

A trivial explanation for these results could be that this correlation results from general similarities across the two behavioral epochs, such as having equivalent tasks or shared layout of the virtual worlds. To probe the structure of inhibition suppression in a drastically different context, we used the “no task” epoch described previously (Figure 4). Here the visual scene is distinct and static, and there is no behavioral task. Even here, when comparing activity suppression in New 1 vs. “no task” epochs, we found significant correlation for each cell on day 1, indicating that the functional inhibitory network structure for these two very different behavioral epochs is very similar. We also measured correlation in activity suppression throughout the 5 day protocol. While the correlation was significant on day 1 (Figure 5B, r=0.46 p<0.001), it was not for days 2 – 5 (Figure 5B, Day 2, r=0.089, p>0.999; Day 3 r=0.026, p>0.999; Day 4 r=0.016, p>0.999; Day 5, r=-0.077, p>0.999). This was not surprising because inhibitory activity in New 1 recovers as animals learn the task, while suppression remains strong with no learning in “no task” epochs.

The rapid time-dependent loss of correlation between New 1 and “no task” epochs reinforces how striking the correlation structure is for New 1 vs. New 2, indicating that not only is initial suppression correlated but there is also consistency in the temporal dynamics of this structure in time across five days (Figure 5C). Similarly, the mean difference in percent change for each cell between New 1 and New 2 remains stable across the remapping paradigm (Spearman’s ρ=0.074, p=0.173), whereas the difference between New 1 and the “no task” epoch steadily increases over the five day course of exposure to each environment (Figure 5D, Spearman’s ρ=0.39, p<0.001).

These results show that each cell exhibits a consistent degree of activity suppression during learning. To understand more about how this structure arises, we looked for other factors that were associated with each cell’s magnitude of activity suppression. First, we examined the recovery of inhibition in cells as a function of initial level of activity suppression on day 1 of remapping into New 1. Cells stratified by the magnitude of their initial suppression continue to be stratified by activity suppression across the five day remapping protocol, with the most strongly suppressed on day 1 remaining the most suppressed on day 5, while the least suppressed remain the least suppressed (Figure 5—figure supplement 1A). This finding indicates that each cell’s initial activity suppression is predictive of future activity throughout the protocol, suggesting that neurons may not be drawn from the same population of functionally homogeneous interneurons. Similarly, cells which were most suppressed in the no task epoch remained the most suppressed through subsequent days of exposure to this epoch (Figure 5— figure supplement 1B).

Does the degree of activity suppression differentially map onto distinct SOM-int cell types? SOM-ints are primarily composed of two functionally and anatomically distinct types, OLM and bistratified interneurons (Klausberger et al., 2004; Royer et al., 2012). The somata of OLM neurons lie in *stratum oriens* (SO), while bistratified interneurons are mostly in *stratum pyramidale* (SP). We found no difference in activity suppression between cells with somata in SO vs. SP (Figure 5—figure supplement 1C, p=0.38). Baseline fluorescence (which may be used, with limitations, as an indicator of basal firing rate) also exhibited no relationship to activity suppression (Figure 5—figure supplement 1D Spearman’s ρ=-0.16 p=0.17).

We previously identified two distinct populations of SOM-ints, one whose activity was positively correlated with locomotion and another whose activity was anti-correlated (Arriaga and Han, 2017). These two populations, as measured by phase angle of the hilbert transform of the cell’s correlation between stop-triggered mean activity and running speed, were also present in these data. However, there was no difference in activity suppression between the two (Figure 5—figure supplement 1F, p>0.99). We also found no relationship between soma area and activity suppression (Figure 5—figure supplement 1E, p>0.99). Thus, the degree of activity suppression was not readily explained by previously identified cell classes or by cellular properties.

Finally, we examined whether activity suppression in New was associated with each cell’s GLM fit of fluorescence. In the cell population, there was variability in the goodness of model fit so we investigated whether model fit in Fam was predictive of activity suppression in New. Using both RMS error and R^2^, we found no relationship between the two variables (Figure 5—figure supplement 1G, H).

## Discussion

In this work we addressed critical questions in neuronal network function: how is inhibition dynamically regulated during learning, and is there a persistent structure in functional inhibitory activity dynamics? We found that the activity of SOM-ints was initially strongly suppressed upon exposure to a novel virtual environment and activity gradually recovered as animals learned the goal-directed spatial navigation task in the second, initially novel, environment. Furthermore, there was a persistent inhibitory network structure in the transition from familiar to novel environments. Each interneuron exhibited a characteristic amount of activity suppression in multiple new environments, as well as in a drastically different “no task” environment where there was no relevant learning.

Our findings are consistent with a model in which a new environment is associated with decreased network inhibition which then gradually recovers over the course of learning to stabilize the network, possibly via synaptic plasticity (Cohen et al., 2017; Wilson and McNaughton, 1993). We tie inhibitory suppression to learning in two ways. First, using learning of a goal-directed spatial navigation task, we were able to produce prolonged suppression of SOM-ints. This prolonged SOM-int suppression overlapped with learning of the new environment, in particular reward location, as evidenced on behavioral measures including task performance, reward-associated locomotion, and licking patterns. Second, we tested the role of learning in recovery of inhibitory activity by providing an epoch with no task, and therefore no learning. We found that inhibition remained suppressed when animals are not engaging in spatial learning.

Two other noteworthy observations arise from this “no task” experiment. First, inhibition was strongly suppressed even though the animal was not learning, suggesting that inhibition suppression is characteristic of a network primed for learning, rather than an indication of novelty. Second, locomotion speed is significantly greater in the “no task” epoch on day 1 than in Fam. Based on the general positive correlation between movement speed and interneuron activity (Arriaga and Han, 2017) we would expect inhibitory activity to be greater in the “no task” epoch, yet it is far lower. This result, in combination with the decoupling of locomotion and activity during learning in the new environment, highlights the importance of behavioral state in controlling the output mode of neurons.

We have shown long-lived SOM-int suppression that is distinct from transient suppression of SOM-ints during place cell global remapping. How much inhibition suppression is associated with novelty-induced place cell global remapping and how much is due to learning *per se?* In both real world and VR experiments, switching animals to new environments triggers a few minutes of inhibition suppression and formation of new place cell maps on the same timescale (Frank et al., 2004; Hainmueller and Bartos, 2018; Muller and Kubie, 1987; Nitz and McNaughton, 2004; Sheffield and Dombeck, 2015; Wilson and McNaughton, 1993). In contrast, in a learning task with no global remapping where freely moving rats learned new goal locations in a familiar environment, fast-spiking putative interneurons both increased and decreased activity as performance increased (Dupret et al., 2013). This reflected dissolution of ensembles encoding old reward maps and formation of new cell assemblies, comprising pyramidal and interneurons, representing new reward locations. Taken together, it is likely that the initial SOM-int activity suppression in our experiments is triggered by the switch into a new context and associated global remapping while slowly increasing inhibition thereafter is associated with additional map refinement related to task-learning. While it is likely that learning shares significant common network mechanisms with place cell remapping, further experiments using more detailed simultaneous recordings of pyramidal and interneuronal activity will refine this picture, while more subtle environmental manipulations will help dissociate global remapping effects from learning.

Another critical contribution of this study was the discovery of stable inhibitory activity dynamics during learning, enabled by our stable, long-term recording of identified interneurons. We found inhibitory population structure, with some SOM-ints strongly suppressed during learning while others were unaffected. Surprisingly we found that these activity dynamics were stable on a cell-by-cell basis, with each interneuron having a consistent level of activity suppression, both across multiple new environments and in a “no task” epoch. These findings reveal an underlying inhibitory circuit structure that is observable when the animal is primed for learning.

Our finding that activity dynamics of interneurons is at least partially independent of excitatory drivers argues against the commonly accepted model where interneurons are simple passive followers of pyramidal activity. This consistent inhibitory framework instead suggests that inhibition shapes the encoding of information in the network by regulating activity in connected ensembles of pyramidal neurons. It is possible that pyramidal neurons downstream of strongly suppressed interneurons are more likely to express plasticity during learning, as a direct result of increased activity due to release of inhibition. Conversely pyramidal neurons downstream of less suppressed interneurons will receive relatively normal levels of inhibition during learning, perhaps limiting plasticity. Thus this functionally diverse, but consistent, inhibitory structure may act as a mechanism to address a fundamental tradeoff in neuronal network function: balancing stability with plasticity (Abraham and Robins, 2005; McClelland et al., 1995; McCloskey and Cohen, 1989). By modulating the functional properties of downstream neurons, inhibition can create plastic and stable pyramidal ensembles, allowing the learning of new information while preserving existing network function. The regulation of specific pyramidal neuron ensembles through interneuron control has not been well-studied (Rao-Ruiz et al., 2019); however, intriguing evidence finding distinct pyramidal neuron populations that code either for learning (engram cells) or stable place coding over learning, support the existence of hippocampal microcircuits that specialize in plasticity or stability (Tanaka et al., 2018).

One requirement of this model is preferential or targeted connectivity in the outputs of SOM-ints onto pyramidal neurons. While such a scenario is at odds with the prevailing view of “pooled” or “blanket” inhibition where interneurons make promiscuous and non-selective synapses (Fino and Yuste, 2011; Packer and Yuste, 2011), significant evidence exists for preferential connectivity both in the hippocampus and cortex. In the hippocampus, PV-expressing basket cells preferentially inhibit deep pyramidal neurons projecting to the amygdala while also being more likely to receive excitation from superficial pyramidal neurons or deep pyramidal neurons projecting to the prefrontal cortex (Lee et al., 2014). In the medial entorhinal cortex, cholecystokinin-expressing basket cells selectively target pyramidal neurons that project extra-hippocampally (Varga et al., 2010). Furthermore, in the hippocampus, interneurons participate in cell assemblies with pyramidal neurons and can share coding properties such as place fields (Ego-Stengel and Wilson, 2007; Kubie et al., 1990; Marshall et al., 2002). Similarly, functional subnetworks of interneurons and pyramidal neurons have been identified in the cortex (Khan et al., 2018; Najafi et al., 2019; Znamenskiy et al., 2018). Finally, this work identifying specialization of interneuron function is complemented by evidence for functionally distinct subsets of CA1 pyramidal neurons (Danielson et al., 2016; Graves et al., 2012; Mizuseki et al., 2011; Soltesz and Losonczy, 2018).

We identified structured activity dynamics in the functional responses of interneurons as animals learned a task in novel virtual environments. What mechanisms might generate the activity dynamics and structure within the interneuron population? Neuromodulatory transmitters targeting G-protein coupled receptors are likely to play a significant role. Novelty or arousal produce strong changes in neuromodulation, with sharp increases in acetylcholine (Acquas et al., 1996; Vinck et al., 2015), norepinephrine (Sara et al., 1995), and dopamine (Kempadoo et al., 2016; McNamara et al., 2014; Takeuchi et al., 2016), among others. Differing levels of inhibitory activity suppression could be set by expression levels of neuromodulatory receptors in each cell. For example, interneurons show markedly divergent responses to acetylcholine depending on their composition and expression of receptors (McQuiston and Madison, 1999).

Another possible mechanism for suppressing inhibition is disinhibitory connections from other interneurons targeting SOM-ints. In the hippocampus, this disinhibitory input can come from local VIP, PV, and SOM interneurons (Francavilla et al., 2015; Lovett-Barron et al., 2012). Indeed, in our experiments some SOM-ints were activated in novel environments, although it remains unclear if these interneurons provide disinhibitory input. VIP interneurons are strongly associated with disinhibition and previous work showed that these neurons are necessary for hippocampal-dependent learning (Donato et al., 2013; Turi et al., 2019).

Finally, it is possible that decreased SOM-int activity is inherited from reduced upstream excitatory input. In this case feed-forward inhibition is driven by EC and CA2/3 input while feed-back excitation is driven by local CA1 neurons. However, during learning or novelty, CA3 and EC pyramidal neurons don’t change firing rates, while CA1 pyramidal neurons increase activity (Barry et al., 2012; Karlsson and Frank, 2008; Nitz and McNaughton, 2004; Wilson and McNaughton, 1993) making it unlikely that inhibitory suppression is purely a function of reduced excitatory drive.

Our work identifies inhibitory activity dynamics that are tied to learning while revealing that individual interneurons have a consistent functional role across learning episodes. Together, this work and others highlight functional specialization within defined sets of neurons which may serve to allow efficient incorporation of new information while maintaining overall network stability.

## Materials and Methods

### Animals

All experiments were approved by the Washington University Animal Care and Use Committee. Heterozygotes (+/−) from two cre-driver mice lines on a C57Bl/6J genetic background were used to label parvalbumin-expressing and somatostatin-expressing inhibitory interneurons: SST^tm2.1(cre)Zjh^/J (SOM-cre) and Pvalb^tm1(cre)Arbr^/J (PV-cre; Jackson Labs). All imaging data were from SOM-ints while behavioral data in Figure 1 were from PV- and SOM-ints.

### Viral Injections and hippocampal window implantation

Surgical procedures, VR track running behavior, and two-photon imaging have been described previously (Arriaga and Han 2017). Briefly, mice were injected with adeno-associated virus (AAV) at 2-4 months of age. Mice were anesthetized with isoflurane, and a small (.5mm) craniotomy was opened above the left cortex. Virus (AAV1.Syn.Flex.GCaMP6f.WPRE.SV40, Penn Vector Core, University of Pennsylvania, 1.71 × 10^13^ genome copies, diluted 1:1–1:4 with PBS, ~50nL total volume) was pressure injected through a beveled micro-pipette targeting CA1 (−1.8 ML, −2.0 AP, −1.3 DV).

After virus injection, mice were water-scheduled for 1-3 weeks and an imaging cannula (2.8 mm diameter) was implanted above the hippocampus by aspirating the overlying cortex. Mice recovered for at least two weeks after surgery before beginning training.

### VR track running behavior

The virtual reality display used a custom-built semi-cylindrical projection screen (1 ft radius) and two rear projectors (Optima 750ST). Projection screen was ~12 inches in front of the mouse and occupied 180° of horizontal, 16° below the horizon and 35° above. The mouse was head-fixed on a spherical Styrofoam treadmill supported on a cushion of air from a 3D printed base which allowed free ball rotation with mouse locomotion. Treadmill movement was tracked using a G400 Logitech mouse configured in LabView (National Instruments). The VR environment was rendered using ViRMEn (Virtual Reality Matlab Engine; Aronov and Tank, 2014). Mice were trained to run to alternating ends of a linear VR track (180 cm) for 2-5 weeks until they consistently achieved target performance (>2 rewards/min for one week). After training, mice were imaged during exposure to a new visual virtual world. Remapping experiments consisted of 7 min behavior in the familiar track (Fam), an instantaneous switch to a novel track of the same length with different visual textures and landmarks (New) for 14 min, and then a return to the familiar (Fam’) for 7 min. This remapping protocol was repeated for 5 successive days with the same, decreasingly novel, New world. In a subset of animals, a second remapping task was also performed. This task was identical to the first with the exception of a different New environment (New 2). Additionally, this same subset of animals was imaged in a “no task” session. This session consisted of 7 min of navigation in the familiar track, 14 min of exposure to a dark screen with no rewards, and a return to the initial familiar environment for 7 min.

### Two-Photon Imaging

Calcium imaging was performed on a Neurolabware laser-scanning two-photon microscope, with the addition of an electric tunable lens (ETL; Optotune, EL-10-30-NIR-LD) and f=-100 mm offset lens to rapidly change axial focal length. We imaged 4-6 axial planes spanning up to 250 μm in the z-axis at a total frame rate of 31Hz, resulting in a per plane sampling rate of 5.2Hz for a 6 plane recording and 7.8Hz for a 4 plane recording. Field of view in *x-y* was 500 × 500μm. Laser power (at 920nm) was ~25 – 50mW after the objective and was set independently for each plane imaged.

### Data Analysis

Data were analyzed using custom programs written in Matlab (RRID:SCR_001622). Images were motion-corrected using cross-correlation registration and rigid translation of individual frames. Slow fluctuations in fluorescence were removed from calculations of ΔF/F_0_ by calculating F_0_ using the eighth percentile of fluorescence intensity from a sliding window 300 s around each time point. ROIs were selected using a semi-automated process. Possible ROIs were identified as contiguous regions with SD>1.5 and an area >90 μm^2^. Overlapping ROIs were manually separated, ROIs were redrawn by hand to separate adjacent cells into distinct ROIs. Unresponsive puncta, or those with low signal-to-noise ratios (initially identified as having a skewness of ΔF/F in the first familiar environment less than 0.3) were dropped from further analysis. When the same cell was recorded in multiple planes, the brightest ROI was used. Neuropil contamination was removed by subtracting a perisomatic fluorescence signal from an annulus between 5 and 20 μm from each ROI, excluding any other possible ROIs (F_Corrected-ROI_, =F_ROI_ –.8 * F_Neuropil_).

The percent change in the New environment was calculated on each day for each cell as the ratio between the mean fluorescence in the 14 min New world exposure and the mean of the fluorescence from the two 7 min familiar worlds exposures, normalized by the sum of these means 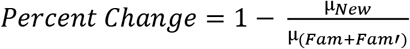.

### Behavior Analysis

Ball movement data, sampled at 1kHz, was downsampled to match the imaging frame rate. All normalized behavioral metrics were normalized by taking the ratio of the metric in the New world to Fam_*Ave*_ (mean of Fam and Fam’). Task performance was calculated as the rewards received per minute. Speed was calculated as the Euclidean sum of forward and rotation components of ball velocity. Deceleration was calculated as the first derivative of the forward component of the ball speed during a three second window prior to reward. Location of trial failure was identified as the distance from the correct destination end zone where the animal turns around before reaching the end zone.

Lick behavior was detected using a 2-transistor lick detection circuit (Slotnick 2009). Individual licks were not resolvable, so lick responses were binned into lick bouts, defined as a period of repeated lick responses with less than 200 ms between repeated lick signals. The lick rate was calculated as the number of these licking bouts per minute. The fraction of “correct” licks was calculated as the fraction of licking bouts which occur within ±500ms of reward delivery (marked by an audible solenoid click to dispense water). The fraction of licks in unrewarded end zones was calculated as the fraction of incorrect, unrewarded, entries into the track end zone which elicited a bout of licking.

### General Linear Model of Activity

A general linear model was used to estimate fluorescence as a function of the behavioral parameters which are correlated with cell activity. The model predicts fluorescence as the linear combination of weighted, time-lagged behavior components. The lag used for each component was determined by the time of the peak of its cross-correlation with cell activity. Modeling of interneuron fluorescence was done using the *glmfit* function in Matlab with a normal distribution and an identity link function. The oscillatory nature of interneuron fluorescent activity series, without the large transients typical in pyramidal cells, were better fit using a normal distribution than the Poisson distribution commonly used in generalized linear models of pyramidal cell activity. Models were trained using fluorescence data from Fam and fit using the forward and rotation components of ball speed, the timing of rewards, and the position and speed in the virtual reality environment. Behavioral data was included at a lag of up to 2 seconds, determined by the maximum value of the cross-correlation between each parameter and cell activity. Root mean square (RMS) error and model correlation in Fam were calculated using 10-fold cross validation, successive models were trained on 9/10 of the data set and tested on 1/10 of the data, the average model performance across these ten sessions was used in Fam. performance in other sessions was calculated by applying the model fits trained in Fam to behavioral data from subsequent epochs. Correlation was measured as the Pearson correlation between modeled traces and ΔF/F in each context.

### Experimental Design and Statistical Analysis

Behavior data is reported from 9 mice (5 male, 4 female). We recorded 162 somatostatin-cre positive cells (mean=20.3+/− 4.9) from eight mice (6 male, 2 female) across all 5 days of the initial remapping experiment. For the second remapping and “no task” paradigms we recorded from 4 of these mice (3 male, 1 female), tracking 69 cells across all 3 contexts.

Significance of normalized data metrics were calculated using 1 sample t-tests of mean values, differences between familiar and new epochs were calculated using paired sample t-tests of mean values, RMS error values were calculated on each cell in each epoch with paired-sample t-tests, Trends over days were assessed using Spearman’s Rank Correlation. Significance of RMS error and R^2^ values were calculated on each cell in each epoch with paired-sample t-tests with Bonferroni-Holm corrections. Correlations of percent change across paradigms were calculated using Pearson Correlation. Pearson correlations were used to calculate the correlation between percent change in each New world or No Task epoch. All multiple comparisons were corrected with Bonferroni-Holm Corrections.

Locomotion and immobility associated interneurons were identified using the phase angle of the cross-correlation of interneuron activity and running speed, as described previously (Arriaga and Han, 2017).

Statistical analyses were performed in Matlab.

## Acknowledgements

This work was supported by the McDonnell Centers for Systems Neuroscience and Cellular and Molecular Neurobiology, and through the Cognitive, Computational, Systems Neuroscience Pathway at Washington University in St. Louis.

---

**Figure 1—figure supplement 1:**
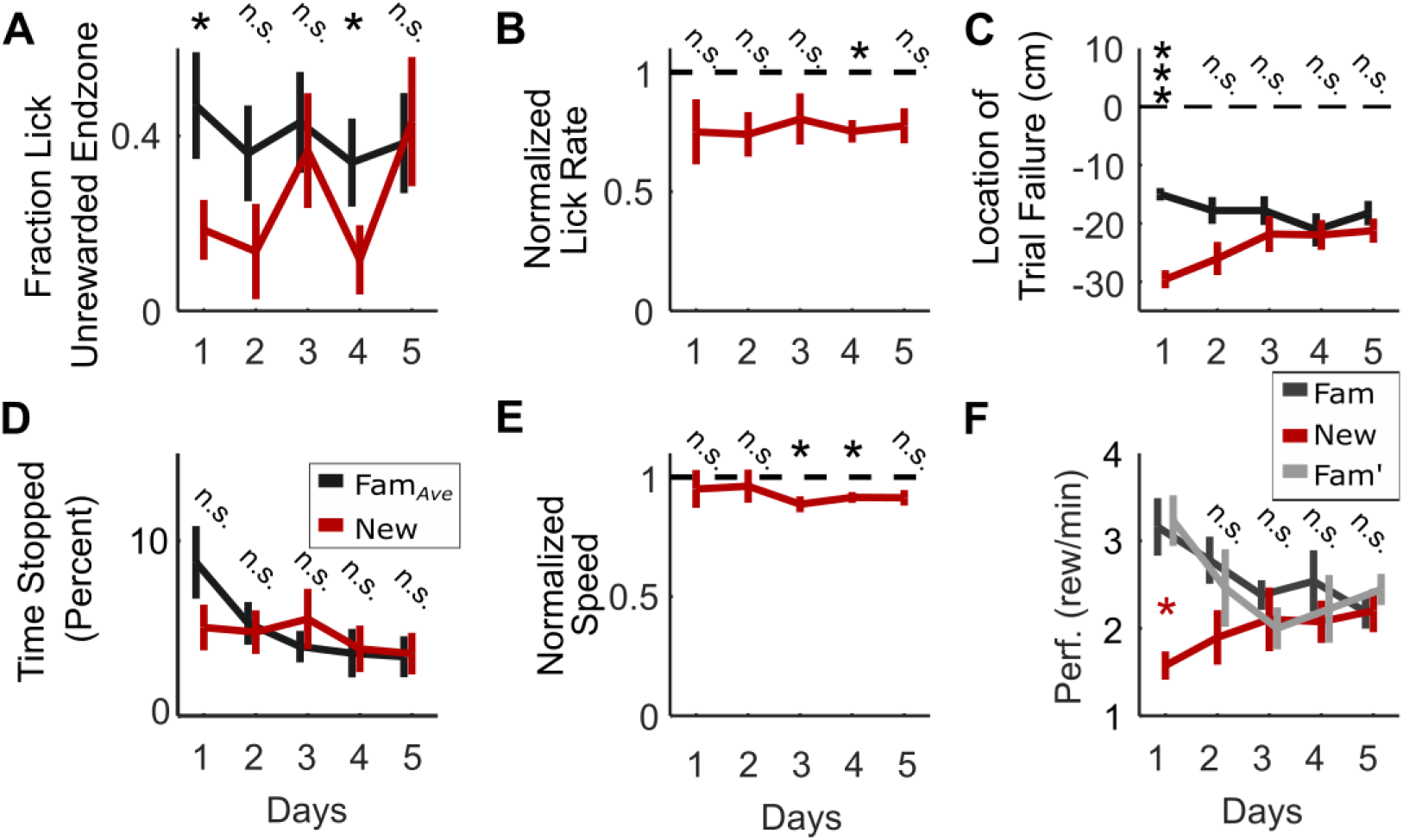
Behavior metrics in New world. **A**, Mice are significantly more likely to lick in expectation of a reward upon incorrectly entering an endzone in the familiar environment (when they return to the same reward zone twice in a row), suggesting they know reward locations in Fam but not New. This difference decreases over the course of exposure to the New world. Fraction of incorrect, unrewarded, entries into a track endzone which coincide with a bout of licking across five days of remapping. **B**, Mice lick at similar rates in both the familiar environment and the New world. Frequency of bouts of licking in new world across five days of remapping, normalized to mean rate of licking in familiar worlds. **C**, Failed trials (where animals turn around too early in the track and return to the same end zone, resulting in no reward) in the New world are farther from correct end zones than in the familiar environment. This difference decreases over 5 days of remapping. Mean distance from endzone of missed trial, identified as the location of a premature turn in the track. **D**, Mice spend similar amount of time stopped in both environments over 5 days, indicating that stopped periods do not significantly contribute to decreased behavioral performance. **E**, Average running speed is similar in both Fam. and the New world, suggesting slower running does not contribute to decreased behavioral performance on day 1. Running speed in New across five days of remapping, normalized to mean speed in Fam_*Ave*_. **F**, Mice were initially impaired in behavioral performance in the new environment relative to both Fam and Fam’. Data show return to high performance in Fam’, suggesting that satiation or fatigue do not contribute to impaired behavioral performance in New. Mean performance, measured in rewards per minute, in New world over 5 days of remapping. (n.s. p>0.05, *p<0.05, **p<.01, ***p<.001 by paired sample t-test or 1-sample t-test with Bonferroni-Holm Corrections, N=9).

**Figure 1—figure supplement 1:**
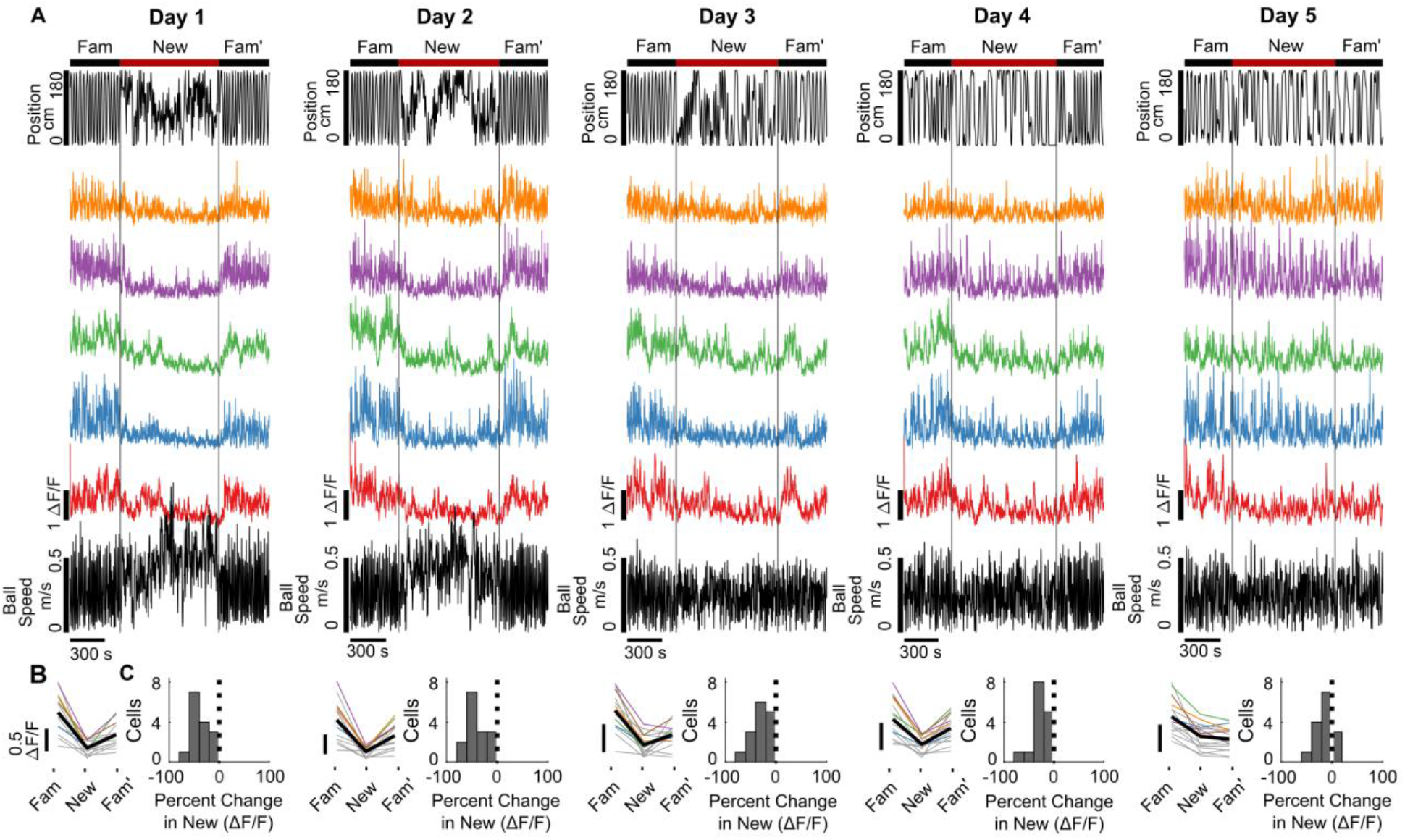
SOM-int activity suppression over five days of remapping into New. **A**, Cellular activity is initially strongly suppressed but recovers over multiple exposures to New. Top, position in VR track, middle, ΔF/F of sample cells, bottom, ball speed. **B**, Mean ΔF/F of all cells from example mouse on day 1 of remapping (colors) and mean (black). **C**, Histogram of percent change in ΔF/F of SOM-ints in New world relative to Fam_*Ave*_ across five days of remapping. (n=25).

**Figure 1—figure supplement 1:**
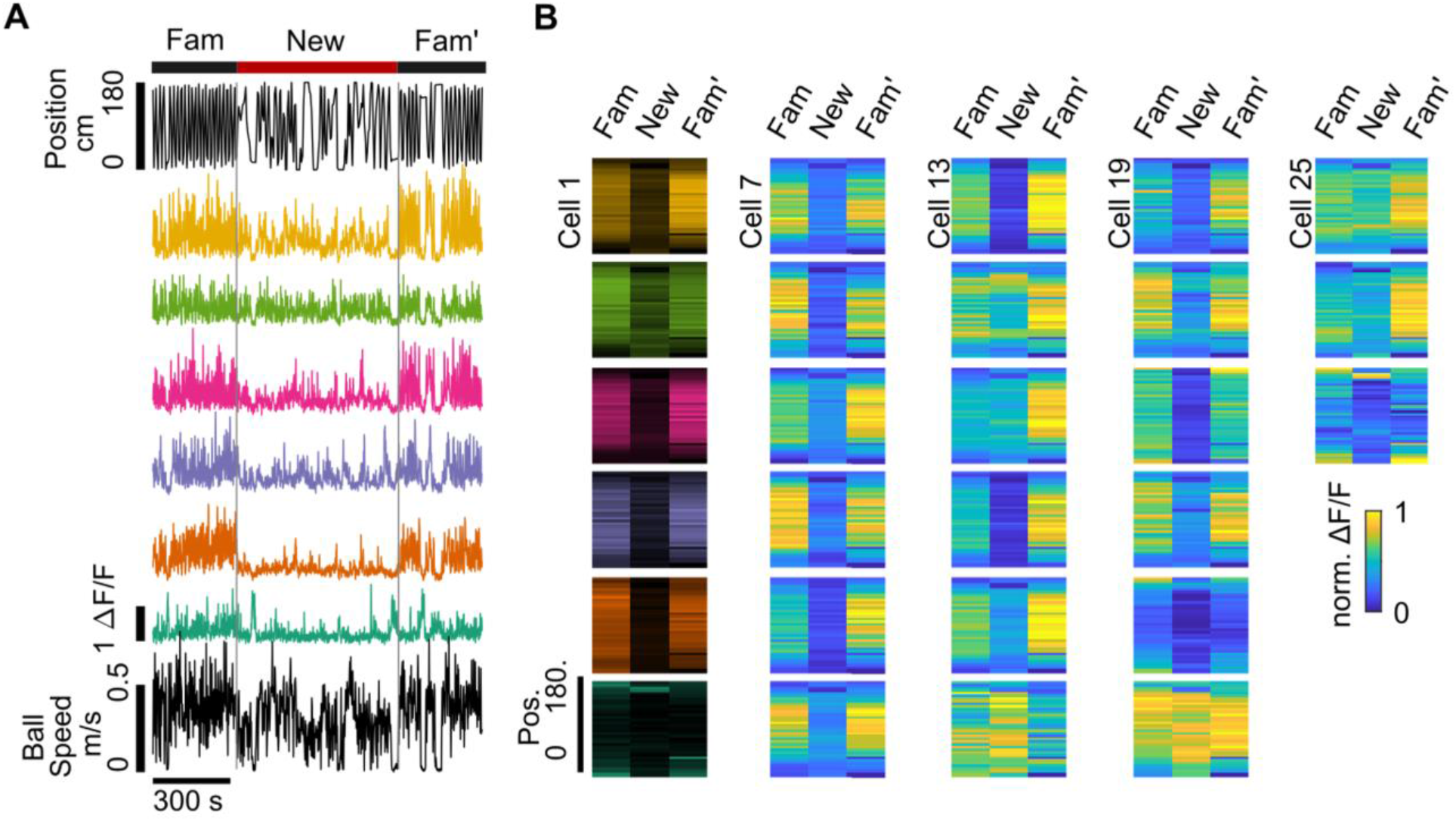
SOM-int firing fields in Fam and New on day 1. Data from same sample mouse in Fig. 2A – C.**A**, Top, position in VR track, middle, ΔF/F of sample cells, bottom, ball speed. **B**, SOM-int firing is broadly tuned in Fam and suppressed in New. Heatmaps of neuronal activity in the VR track on day 1 of remapping for the 28 cells in this example mouse. Cells 1 – 6 are the cells shown in (A), with the same color of heatmap.

**Figure 3—figure supplement 1:**
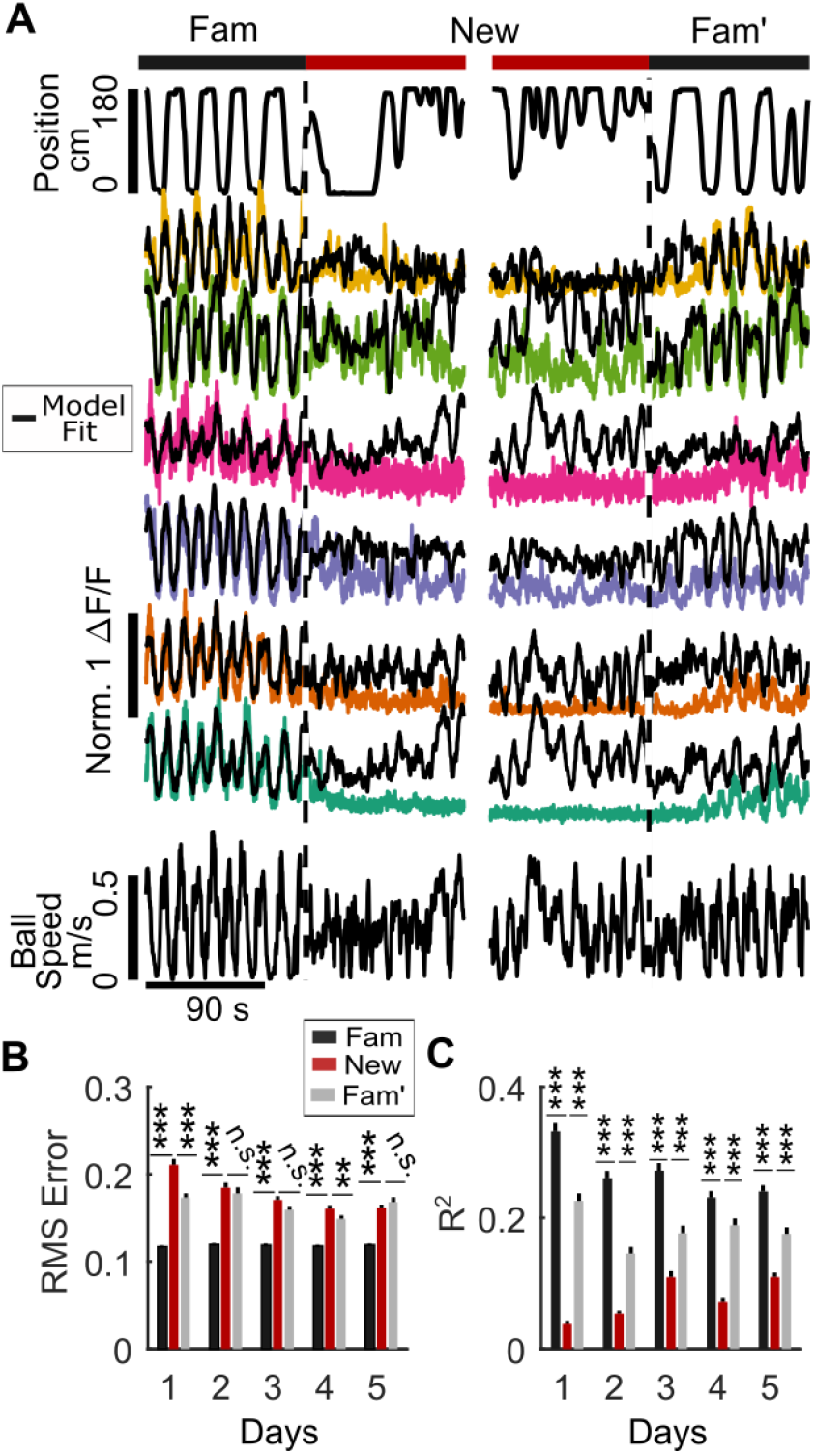
GLM performance in different environments. **A**, On day 1 in New, modeled ΔF/F (black) is larger than actual ΔF/F (colored traces), while in Fam’, modeled fit improves relative to New. **B**, RMS error of model fit is significantly different in Fam versus New on all days, while New is different from Fam’ on day 1 and 4. **C**, Average R^2^ between modeled fluorescence and cell fluorescence across environments and days. (*p<0.05, **p<.01, ***p<.001 by paired sample t-test with Bonferroni-Holm Corrections N=8, n=162).

**Figure 3—figure supplement 2:**
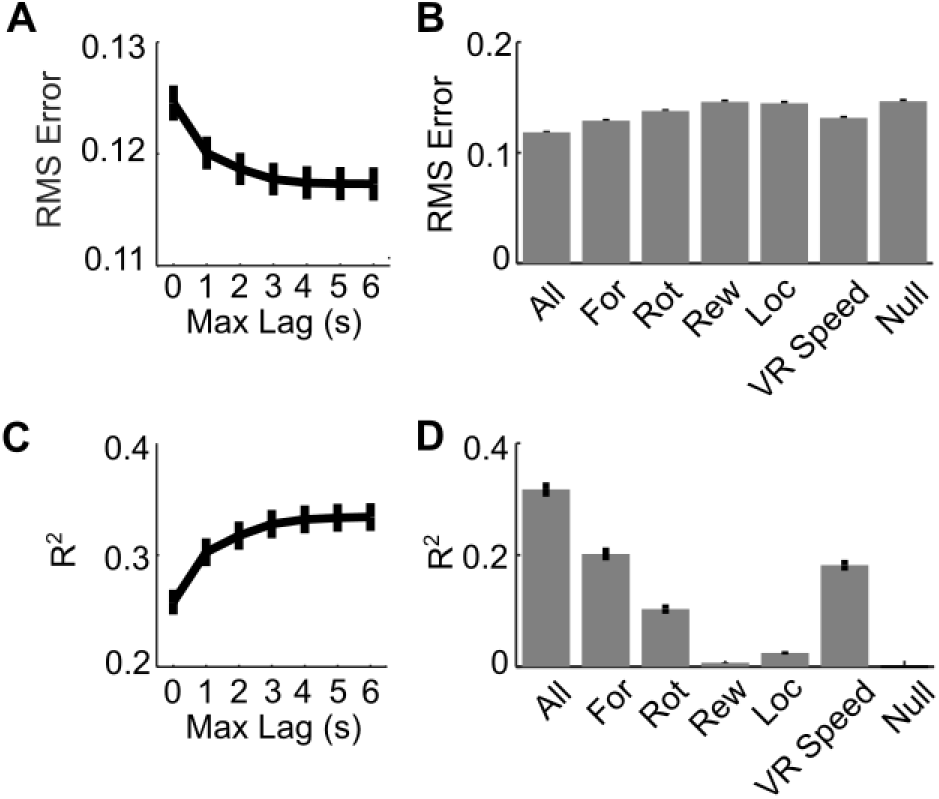
Features which influence model fit. **A**, Using behavioral data at increasing possible lag values improves model performance. Linear models were trained on behavioral data with varying amount of time permitted in the range used to identify the peak of the cross correlation between cell activity and behavioral parameters. Model error (root mean square) decreases with amount of lag included in the model. **B**, Linear models were trained using only one of the parameters used to train the full model to examine the relative importance of different parameters to model performance. Model error (2 seconds lag used) is lowest when including all features used to train model. Relative performance of model trained on only one feature varies. For: Forward component of running speed, Rot: Rotation component of running speed, Rew: Timing of rewards, Loc: Position in VR track, VR Speed: Speed in virtual reality environment, Null: constant model at mean firing rate. **C**, Percent of variance explained by model (R^2^) increases with amount of lag included in the model. **D**, Percent of variance (R^2^) explained by model (2 seconds lag used) is highest when including all features used to train model. (N=8, n=162).

**Figure 4—figure supplement 1:**
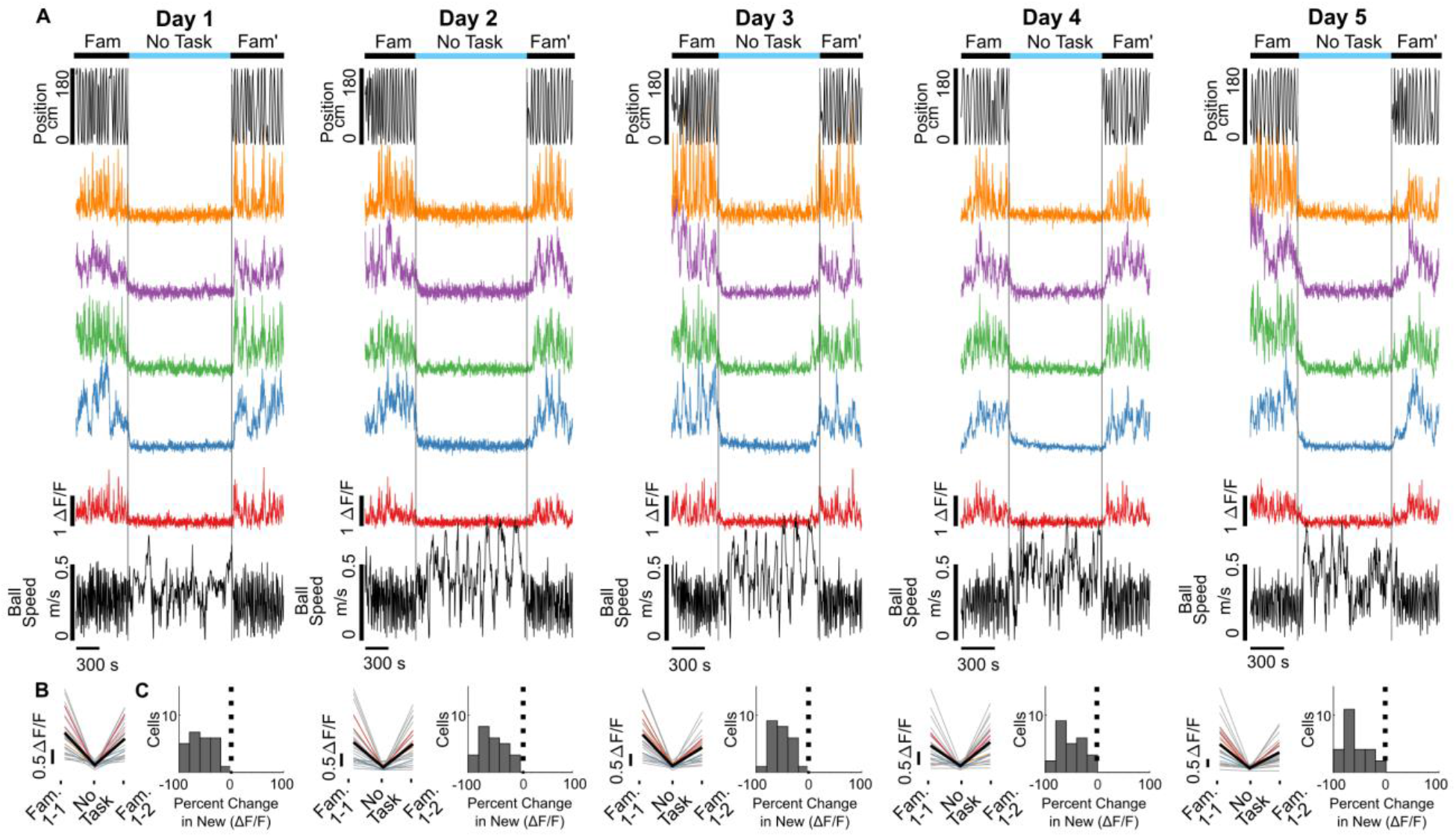
SOM-int activity suppression in “no task” environment. Cellular activity is remains strongly suppressed over multiple exposures to “no task” environment. **A**, Top, position in VR track, middle, ΔF/F of sample cells, bottom, ball speed. **B**, Mean ΔF/F of all cells from example mouse on day 1 of “no task” environment. **C**, Histogram of percent change in ΔF/F of SOM-ints in “no task” relative to Fam_*Ave*_ across five days of exposure. (n=24).

**Figure 5—figure supplement 1:**
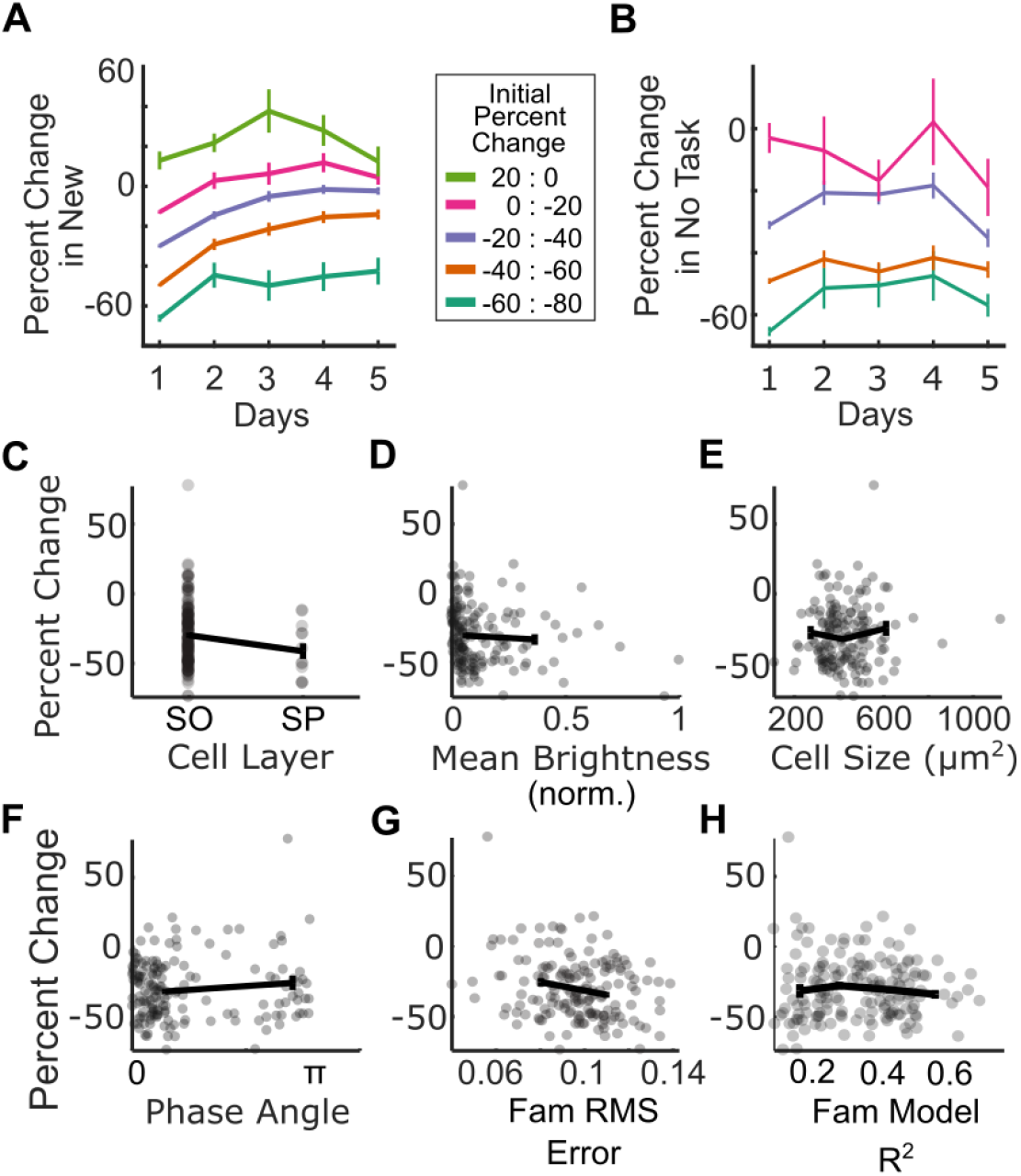
Characterization of suppressed interneurons. **A**, SOM-ints most inhibited on day 1 in New remain the most suppressed over the course of the experiment. Percent change in activity in New relative to Fam_*Ave*_, across five days, stratified by degree of suppression on day 1. **B**, Percent change in activity in “no task” session, stratified by degree of suppression on day 1. **C**, Soma location was not associated with inhibition suppression in New. Percent change of activity on day 1 in New based on soma location, either stratum oriens (SO) or stratum pyramidale (SP). Most SOM-expressing bistratified interneurons have somas in SP while most OLM interneurons have somas in SO. **D**, Mean cell brightness was not associated of inhibition suppression in New. Percent change of activity on day 1 in New versus mean cell brightness. **E**, Soma cell size was not associated with inhibition suppression in New. Percent change of activity on day 1 of New versus cell size. **F**, SOM-int activity correlation with locomotion was not associated of inhibition suppression in New. Percent change of activity on day 1 in New versus phase angle of the Hilbert transform of each cell’s correlation between stop-triggered mean activity and running speed. Cells with a positive activity correlation with locomotion have a phase angle near 0, while those that are anti-correlated are shifted ~180° or π radians. **G**, GLM model fit was not associated of inhibition suppression in New. Percent change of activity on day 1 in New versus RMS error of model fit to actual cellular ΔF/F in Fam. **H**, Percent change of activity on day 1 in New versus R^2^ of model fit to actual cellular ΔF/F in Fam. (N=8, n=162).

